# Volatiles defenses of Amazon *Azteca* ants (repellent ants)

**DOI:** 10.1101/2020.04.15.043547

**Authors:** Kolby J. Jardine, Luani R. de O. Piva, Tayana B. Rodrigues, Gustavo C. Spanner, Jardel R. Rodrigues, Valdiek S. Menezes, Israel Sampaio, Daniela C. Oliveira, Brunon O. Gimenez, Niro Higuchi, Jeffrey Chambers

## Abstract

*Azteca* ants are widely distributed in the neotropics and have been utilized as natural insect repellent for centuries. *Azteca* oils provide natural defense against herbivores in mutualistic interactions between ants and their host trees. While chemical characterization of oil secretions revealed a composition dominated by iridoids and ketones, the volatile emissions from *Azteca* ants, and therefore the active gas-phase semiochemicals, remain uncharacterized. In this study, we determined the composition of volatile emissions from a sample of an *Azteca* ant nest near the Rio Negro in the central Amazon. We found *Azteca* emissions were composed of a blend of methyl cyclopentyl and methyl cyclopentenyl based volatiles previously identified in *Azteca* oil extracts as potent alarm pheromones. The ketone 6-methyl-5-hepten-2-one, which also waspreviously identified as a major component of the *Azteca* oil, was found to be the dominant volatile emitted. For the first time, we report emissions of the highly volatile ketones 2,3-butadione and acetoin from the *Azteca* nest. Our study has important implications for the better understanding of the ecology and defense strategies of *Azteca* ants in herbivore defense and provides a base for future commercial applications involving *Azteca* ant essential oils as natural insect repellents.

## INTRODUCTION

*Azteca* ants are an arboreal genus of ants of the Dolichoderinae subfamily with 130 described species exclusive to the Neotropics (Shattuck, 1995), often found in cavities of the branches and trunks of common tropical trees like *Cecropia* spp. and *Cordia* spp. (Janzen, 1969; Gianoli et al., 2008) (**Figure 1**). A symbiotic relationship among the host tree is established against invading herbivores through a biological defense as they swarm out of their nest and attack any invaders (Agrawal, 1998; Do Nascimento et al., 1998). In addition, *Azteca* ants possess a potent suite of chemical defenses that effectively repel invaders. Numerous chemical defense compounds are produced by *Azteca* ants resulting in repulsion of invaders and alarm behavior in worker ants (Hölldobler & Wilson, 1990; Agrawal & Dubin-Thaler, 1999). These natural insect repellent properties have long been exploited by natives in the Amazon forest to protect themselves from mosquitoes and other attacking insects; after *Azteca* oil is rubbed on the body, the strong smell repels mosquitoes.

**Fig. 1.**
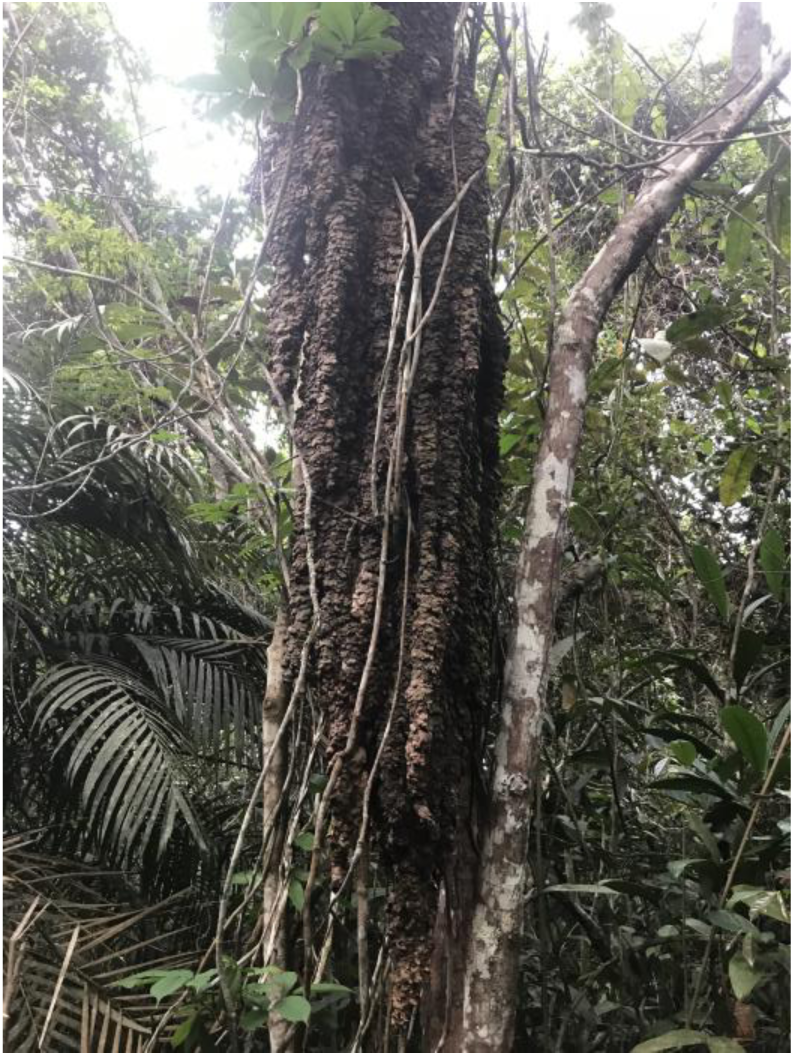
Image of the ant nest where the volatile samples were collected in the central Amazon near the Rio Negro, ∼60 km west of Manaus, Brazil.

*Azteca* oil has not only been used on the human body, but also as a repellent in forest camps or in homes by burning pieces of the flammable ant nest (Posey, 1986). The heat of combustion aids in the volatilization of the oils, thereby helping to clear all of the nearby mosquitoes. These herbivore repellent properties have also been exploited by Kayapó Indians as biological control against pests in agricultural activities (Overall & Posey, 1988; Vandermeer et al., 2002).

The chemical defense agents produced by *Azteca* ants are mediated by volatile organic compounds (VOCs) produced by exocrine glands and excreted as a volatile oil (Attygalle & Morgan, 1984). Chemical analysis of solvent extractions of *Azteca* ant oil have releveled a mixture of iridoids (iridodial isomers, iridomyrmecin, *trans*-*cis*-iridodial, *cis*-*trans*-iridodial and *trans*-*trans*-iridodial, nepetalactone and isodihydronepetalactone) and ketones (2-acetyl-3-methylcyclopentene, *cis*-1-acety1-1-methylcyclopentane, 2-methylcyclopentanone, 2-methyl 1-4-heptanone, 6-methyl-5-hepten-2-one) (Wheeler et al., 1975; Do Nascimento et al., 1998). While the chemical mechanisms of defense deserve additional research, the iridoids have been suggested to be involved in herbivore repulsion while the ketones, including cyclopentyl and cyclopentenyl derivatives seem to be involved in the elicitation of alarm behavior of worker ants (Wheeler et al., 1975; Hölldobler & Wilson, 1990).

Although a chemical characterization of *Azteca* ant oil suggests a composition dominated by iridoids and ketones, the volatile emissions from the oils remains uncharacterized. Emission rates depend both on the concentration of the defense pheromones and herbivore repellants within the oil, and their respective vapor pressures under the ambient temperature conditions. In this study, we characterized the volatile emissions from an intact sample of an *Azteca* ant colony near the Rio Negro in the central Amazon, ∼60 km upstream from Manaus, Brazil. We then compared the composition of the volatile emissions from the nest sample with previously published work on the *Azteca* ant oil composition and discuss the results in terms of herbivore defense mechanisms and potential commercial applications involving *Azteca* ant essential oils as natural insect repellents.

## RESULTS

When the sample of the ant colony was removed with rubber gloves and placed in the chamber for volatile collections on thermal desorption tubes, hundreds of *Azteca* ants swarmed out of the damaged nest sample and entered the chamber together with the nest sample. A strong sweet smell in the surrounding atmosphere could be detected by the researchers conducting the experiments. When compared with the empty glass enclosure, 8 major volatile emissions could be detected from the *Azteca* ant nest sample (see GG-MS selected ion chromatograms in **Figures 2-4**).

**Fig 2.**
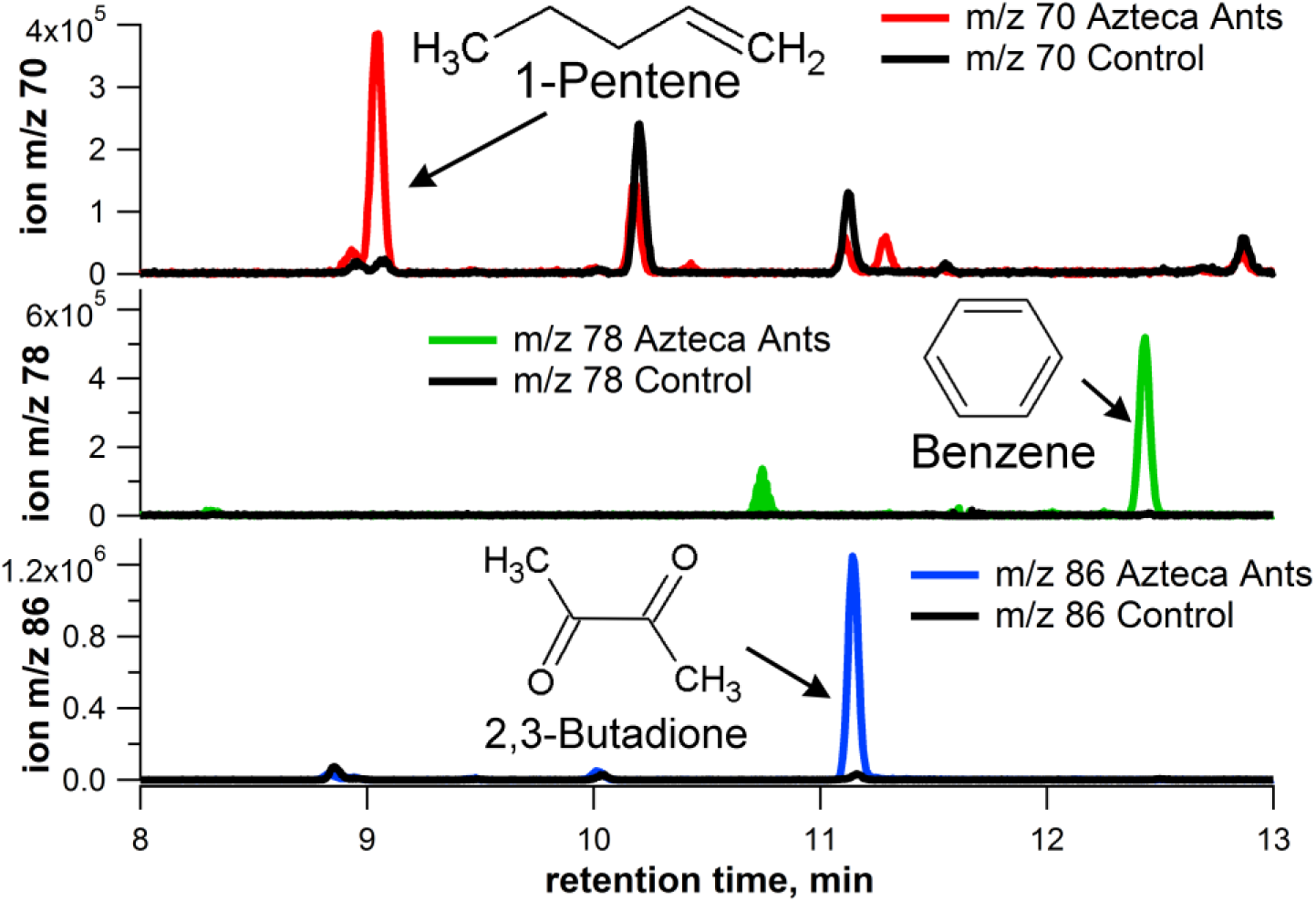
TD-GC-MS selected ion chromatogram of headspace air samples with ants and without (control) showing the presence of 1-pentene, benzene, and 2,3-butadione as emissions from the ant nest.

**Fig. 3.**
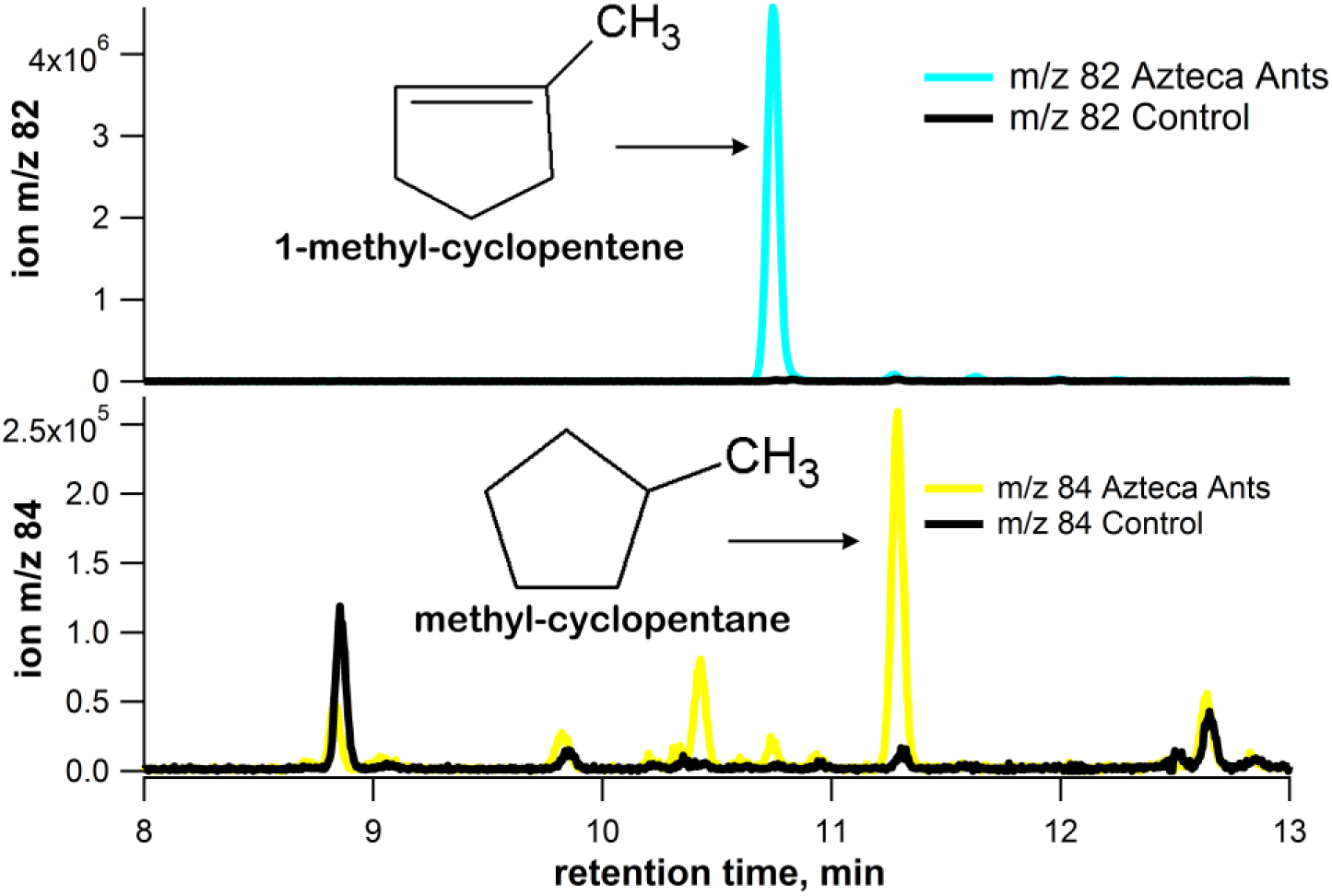
TD-GC-MS selected ion chromatogram of headspace air samples with ants and without (control) showing the presence of 1-methylcyclopentene and methylcyclopentane as emissions from the ant nest.

**Fig. 4.**
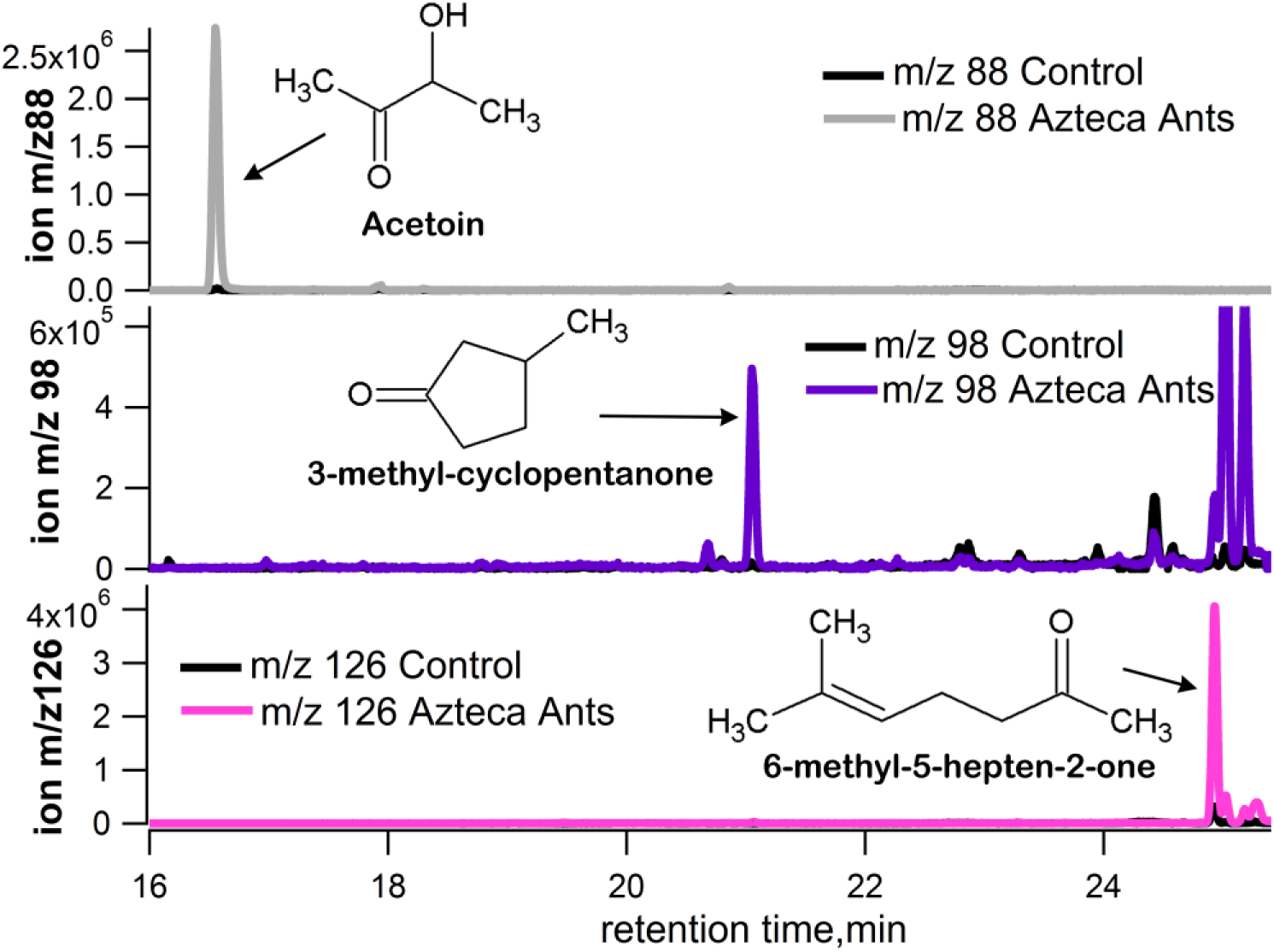
TD-GC-MS selected ion chromatogram of headspace air samples with ants and without (control) showing the presence of Acetoin, 3-methylcyclopentanone, and 6-methyl-5-hepten-2-one, as emissions from the ant nest.

The first group of compounds shared a common structure with a five-carbon ring and included methylcyclopentane, 1-methylcyclopentene, and 3-methylcyclopentanone (Fig. 3-4). In addition, emissions of an acyclic 5-carbon compound (1-pentene) could also be detected (Fig. 2). This suggests that *Azteca* ant emissions consist of a group of compounds with a methylcyclopentyl, methylcyclopentenyl, and methylcyclopentanone structure, potentially derived from the same biochemical pathway. In addition, a group of acylic ketones were observed as emissions from the *Azteca* ant nest including 2,3-butadione, acetoin, and 6-methyl-5-hepten-2-one (Fig. 2 and Fig. 4). Moreover, clear emissions of benzene were observed from the *Azteca* ant nest as well as acetone. However, acetone showed substantial background peaks in the empty enclosure, likely reflecting its abundance in the ambient air (data not shown).

When the composition of the emissions was quantitatively evaluated based on the total ion chromatogram, emissions were dominated by 6-methyl-5-hepten-2-one, which represented 60-70% of total ion peak area. Acetoin and 1-methyl-cyclopentene also represented a major fraction of total emissions from the *Azteca* ant nest with 10-15% of total ion peak area. Emissions of the other compounds were relatively small representing less than 5% of total ion peak area. Emissions of benzene were negligible with less than 0.6% of total ion peak area.

## DISCUSSION

Plant-herbivore interactions drive the coevolution between plants and herbivores, and are mediated, in part, through the emission of VOCs, as direct and indirect defense mechanisms (Kessler & Baldwin, 2001; Fürstenberg-Hägg et al., 2013). VOC emissions may have toxic and repellent effects on herbivores. For example, when under attack by herbivores like beetles, some trees synthesize organic compounds that inhibit the interaction of herbivores to their own pheromones, impairing the efficiency of the attack (Reddy & Guerrero, 2004). Moreover, volatile semiochemicals can be emitted as chemical defenses by insects like *Azteca* ants that interact mutualistic with trees. This chemical defense works synergistically with *Azteca* ant physical defenses, thereby decreasing the chances of a successful attack by herbivores. To date, only qualitative studies evaluating the chemical composition of *Azteca* ant oils have been reported, leaving great uncertainty regarding the composition of gas-phase emissions from the ants and their nest. Therefore, little is known regarding the identity of the major active gas-phase pheromones and repellants emitted within the nest atmosphere and the surrounding atmosphere following an herbivore disturbance.

Consistent with a previous study that identified methyl cyclopentyl and methyl cyclopentenyl compounds as functional alert pheromones in *Azteca* oil secretions [8], in the present study, we observed clear emissions of methyl cyclopentane, 1-methylcyclopentene, and 3-methylcyclopentanone (Fig. 2-4). Wheeler et al. (1975) identified 2-methylcyclopentanone, cis-1-acetyl-2-methylcyclopentane and 2-acetyl-3-methylcyclopentene in the *Azteca* oil. These results suggest that *Azteca* ants are capable of producing a variety of volatile compounds based on methylcyclopentane, methylcyclepentene, and methylcyclopentanone base structures. As 1-methyl-cyclopentene emissions represented an estimated 10-15% of total volatile emissions from the nest, the results suggest it potentially plays an active role as a pheromone in *Azteca* ant alarm signaling.

A group of acylic ketones were observed as emissions from the *Azteca* ant nest including 2,3-butadione, acetoin, and 6-methyl-5-hepten-2-one. We found that volatile emissions from the *Azteca* ant nest sample were dominated by 6-methyl-5-hepten-2-one (representing an estimated 60-70% of total volatile emissions), a ketone previously identified as a major component of *Azteca* ant oil (Janzen, 1969) and potentially elicits alarm behavior in worker ants (Do Nascimento et al., 1998). For the first time, we report the emission of 2,3-butadione and acetoin from an *Azteca* ant nest. These highly volatile ketones were not reported in previous studies on the chemical composition of *Azteca* ant oil. This is likely do to the loss of these highly volatiles ketones during oil extraction and analysis. Our method based on quantitative volatile collection onto thermal desorption tubes is well suited for future studies on the potential pheromone signaling proprieties of these C4 ketones that may be interconverted via redox reactions mediated by a dehydrogenase enzyme (e.g. 2,3-butadione ⟵⟶ acetoin).

The volatile oils excreted from the glands of *Azteca* ants and their gas-phase emissions into the surrounding atmosphere effectively repel invading herbivores both through direct (repellent) and indirect (alarm pheromone) chemical defenses. As trees of the genus *Cecropia* are widespread in the neotropics, their success may be in part due to their mutualistic associations with *Azteca* ants, which can be viewed as an inducible defense for *Cecropia* trees (Agrawal & Dubin-Thaler, 1999). As *Cecropia* are key pioneer species, enabling the rapid recovery of disturbed forests, this suggests a fundamental importance of plant-insect interaction in the ecological maintenance and forest dynamics of tropical Rainforests in the Neotropics.

Although limited information exists regarding the chemical composition of *Azteca* ant oil excretions, information on composition of emissions from the ant and the ant nest hasbeen given little attention. Therefore, the active gas-phase defense and semiochemical agents of *Azteca* ant excretions remain uncharacterized. Future studies should consider that the chemical composition of volatile pheromones and defense compounds of *Azteca* ants may be dependent upon ant and tree species in mutualistic associations, environmental conditions, and specific herbivores involved.

## CONCLUSION

In this study, we verified that several previously identified pheromones present in *Azteca* ant oil excretions are also emitted as gases into the surrounding atmosphere of a damaged nest in the central Amazon forest. This includes 6-methyl-5-hepten-2-one that dominated the emission composition as well as a number of compounds based on methylcyclopentyl and methylcyclopentenyl structures. Future work should verify the source of the emissions by analyzing the volatiles emitted by the ants separately from the ant nest. We report the first 2,3-butadione and acetoin emissions from an *Azteca* ant nest, which represented a significant fraction of total volatile emissions. We speculate that these two VOCs derive from the same biochemical pathway and are interconverted via a redox reaction, likely mediated by a specific enzyme. Future work should verify the C4 ketones as components of *Azteca* ant excretions and evaluate their potential roles as pheromones in herbivore defense responses. Lastly, as oil excretions and volatiles emitted from *Azteca* ants have been widely used by native Brazilians as insect repellants for both personal and agricultural activities, our study provides a base for future commercial applications involving *Azteca* ant essential oils as natural insect repellents.

## EXPERIMENTAL PROCEDURES

An *Azteca* ant nest roughly 2-3 meters in height (1-2 m above the ground) was located near the community of Jaraqui Bela Vista, Brazil (3°00’09” S and 60°24’28” W) in the central Amazon near the Rio Negro (**Figure 1**). A small sample (20 g) of an *Azteca* ant colony was removed from the bottom of one of the nest columns with rubber gloves and immediately placed in a 500 mL glass chamber. Volatile emissions from the *Azteca* ant nest were collected by passing 100 ml min^-1^ of air inside the glass chamber through commercial thermal desorption tubes for 10 minutes (1.0 L air samples). The thermal desorption tubes were commercially purchased and packed with Quartz Wool, Tenax TA, and Carbograph 5TD adsorbents (QTC, Markes International).

The tubes were then transported to the laboratory located in Manaus, Brazil, and within 1 day analyzed for adsorbed volatiles using a thermal desorption-gas chromatograph-mass spectrometer (TD-GC-MS) installed at the National Center for Amazon Research (INPA), as previously described (Jardine et al., 2015). As ambient air entered the chamber through an inlet port to replace air removed from the chamber for volatile collection, background thermal desorption tubes were collected from an empty glass chamber. This was done prior to placing the ant nest sample inside the enclosure in order to test the analytical background of the chamber for each compound analyzed. Volatiles were identified based on mass spectral comparisons with the 2011 National Institute of Standards and Technology mass spectral database. Identification was based on the best match (> 90 % confidence).

## ACKNOWLEDGMENTS

This material is based upon work supported as part of the Next Generation Ecosystem Experiments-Tropics funded by the U.S. Department of Energy, Office of Science, Office of Biological and Environmental Research through contract No. DE-AC02-05CH11231 to LBNL, as part of DOE’s Terrestrial Ecosystem Science Program. Additional funding for this research was provided by the Brazilian Conselho Nacional de Desenvolvimento Científico e Tecnológico. We would like to thank Daniram Pooran at the Anavilhanas Village Hotel (Bela Vista, Brazil) for helping to locate the *Azteca* ant nest.

## AUTHOR CONTRIBUTIONS

K.J. conceived and designed the experiments; K.J., and D.O. performed the experiments; K.J., D.O., T.R., V.M., B.G., L.P., J.C., I.S., G.S., N.H., J.C., analyzed the data, provided logistics support and wrote the paper.

## REFERENCES

Agrawal, A. A. (1998). Leaf damage and associated cues induce aggressive ant recruitment in a neotropical ant-plant. Ecology, 79, 2100–2112.

Agrawal, A. A., & Dubin-Thaler, B. J. (1999). Induced responses to herbivory in the Neotropical ant-plant association between ants and *Cecropia* trees: response of ants to potential inducing cues. Behavioral Ecology and Sociobiology, 45, 47–54.

Attygalle, A. B., & Morgan, E. D. (1984). Chemicals from the glands of ants. Chemical Society Reviews, 13, 245–278.

Do Nascimento, R. R., Billen, J., Sant’ana, A. E. G., Morgan, E. D., & Harada, A. Y. (1998). Pygidial gland of NR. *bicolor* and *chartifex*: Morphology and chemical identification of volatile components. Journal of chemical ecology, 24, 1629–1637.

Fürstenberg-Hägg, J., Zagrobelny, M., & Bak, S. (2013). Plant defense against insect herbivores. International journal of molecular sciences, 14, 10242–10297.

Gianoli, E., Sendoya, S., Vargas, F., Mejía, P., Jaffe, R., Rodríguez, M., et al. (2008). Patterns of ants’ defence of *Cecropia* trees in a tropical rainforest: support for optimal defence theory. Ecological Research, 23, 905–908.

Hölldobler, B., & Wilson, E. O. (1990). The ants. Springer, Berlin: Harvard University Press, 732p.

Janzen, D. H. (1969). Allelopathy by myrmecophytes: the ant as an allelopathic agent of *Cecropia*. Ecology, 50, 147–153.

Jardine, A., Jardine, K., Fuentes, J., Martin, S., Martins, G., & Durgante, F., et al. (2015). Highly reactive light-dependent monoterpenes in the Amazon. Geophysical Research Letters, 42, 1576–1583.

Kessler, A., & Baldwin, I. T. (2001). Defensive function of herbivore-induced plant volatile emissions in nature. Science, 291, 2141–2144.

Overall, W., & Posey, D. A. (1990). Uso de formigas spp. para controle biológico de pragas agrícolas entre os índios Kayapó do Brasil, In D. A. Posey & W. L. Overal (Eds.), Ethnobiology : implications and applications, 2, (pp. 219–25).

Posey, D. A. (1986). Etnoentomologia de tribos indígenas da Amazônia. Suma etnológica brasileira, 1, 251–271.

Reddy, G. V., & Guerrero, A. (2004). Interactions of insect pheromones and plant semiochemicals. Trends in plant science, 9, 253–261.

Shattuck, S. O. (1995). Generic-level relationships within the ant subfamily Dolichoderinae (Hymenoptera: Formicidae). Systematic Entomology, 20, 217–228.

Vandermeer, J., Perfecto, I., Nuñez, G. I., Phillpott, S., & Ballinas, A. G. (2002). Ants (sp.) as potential biological control agents in shade coffee production in Chiapas, Mexico. Agroforestry systems, 56, 271–276.

Wheeler, J. W., Evans, S. L., Blum, M. S., & Torgerson, R. L. (1975). Cyclopentyl ketones: identification and function in ants. Science, 187, 254–255.

